# Ancestral neural circuits potentiate the origin of a female sexual behavior

**DOI:** 10.1101/2023.12.05.570174

**Authors:** Minhao Li, Dawn S. Chen, Ian P. Junker, Fabianna Szorenyi, Guan Hao Chen, Arnold J. Berger, Aaron A. Comeault, Daniel R. Matute, Yun Ding

**Affiliations:** Department of Biology, University of Pennsylvania, Philadelphia, PA, USA; Department of Biology, University of North Carolina, Chapel Hill, NC, USA

## Abstract

Courtship interactions are remarkably diverse in form and complexity among species. How neural circuits evolve to encode new behaviors that are functionally integrated into these dynamic social interactions is unknown. Here we report a recently originated female sexual behavior in the island endemic *Drosophila* species *D. santomea*, where females signal receptivity to male courtship songs by spreading their wings, which in turn promotes prolonged songs in courting males. Copulation success depends on this female signal and correlates with males’ ability to adjust his singing in such a social feedback loop. Functional comparison of sexual circuitry across species suggests that a pair of descending neurons, which integrates male song stimuli and female internal state to control a conserved female abdominal behavior, drives wing spreading in *D. santomea*. This co-option occurred through the refinement of a pre-existing, plastic circuit that can be optogenetically activated in an outgroup species. Combined, our results show that the ancestral potential of a socially-tuned key circuit node to engage the wing motor program facilitates the expression of a new female behavior in appropriate sensory and motivational contexts. More broadly, our work provides insights into the evolution of social behaviors, particularly female behaviors, and the underlying neural mechanisms.

## Introduction

Social interactions between the sexes during mating are pivotal for their reproductive success^1–5^, and animals often employ a suite of behaviors to communicate their quality and interests to potential mates^5–8^. To maintain reproductive barriers between species while permitting sexual selection within species, courtship interactions are often rapidly diversifying. Courtship behaviors exhibit exceptional diversity in complexity and form, often with quantitative and qualitative differences among even closely-related lineages^8–11^. The real-time production of social behaviors requires complex neural orchestration that integrates external and internal cues to guide adaptive motor responses in relevant social contexts. During the elaboration and diversification of courtship behaviors, how new behaviors are incorporated into existing complex social contexts and neural circuitry in a temporally coordinated and meaningful manner remains unknown.

Newly originated behaviors offer a favorable time window to infer the ancestral and derived states, and to pinpoint the initial changes at play before extensive secondary evolutionary changes mask their origins. However, a system to investigate recently originated social behaviors in species amenable to functional comparison of neural circuits has been lacking. In this study, we leveraged *Drosophila* species as an emerging neural comparative model^12–16^ and established a comparative paradigm to explore the origin of new social behaviors at both behavioral and neural levels. Shifting away from the traditional spotlight on male sexual behaviors^17^, we report a recently originated female behavior in *D. santomea*, referred to as wing spreading (WS), in which a female extends her wings in response to a male’s courtship song to signal her receptivity. Combining a phylogenetic survey, behavioral characterization, and functional manipulation of neural circuits between species, we provide insights into the ultimate and proximate mechanisms underlying the origin of WS. We demonstrate that WS evolved as a new receptive female signal that dynamically shapes a male’s courtship efforts and copulation outcome. We further show that the origin of WS is mediated by the co-option of a descending circuit node that drives a conserved abdominal behavior and the refinement of a latent and plastic ancestral circuit.

## Results

### WS is a newly originated receptive female response to male song

In *Drosophila*, the two sexes typically engage in an extended period of courtship interaction, where a female assesses a male based on his signals such as song, dance, and sex pheromone to inform her copulation decision^2,18^. Females communicate sexual interests through two conserved female-specific displays: vaginal plate opening (VPO), indicative of receptivity^19^, and ovipositor extrusion (OE), indicative of rejection^20,21^. In *D. santomea*, a closely-related species of *D. melanogaster*, we observed that a female may extend her wings laterally when a male vibrates one wing to sing a courtship song (Fig. 1a, Video 1). In response to the female’s wing extension, the male may continue singing in place or approach the female to lick her genitalia, the latter of which may be followed by a copulation attempt. This interaction often occurred repeatedly before copulation (Fig. 1a and Extended Data Fig. 1). The female wing extension behavior has not been reported within the *melanogaster* subgroup but is reminiscent of the female WS behaviors described before copulation in species of some distantly related lineages such as the *virilis* group^19,22,23^. Therefore, we refer to this behavior in *D. santomea* also as WS based on similarities of their motor pattern and the pre-copulatory context, noting that the precise social conditions and functions of WS may differ among species.

**Fig. 1:**
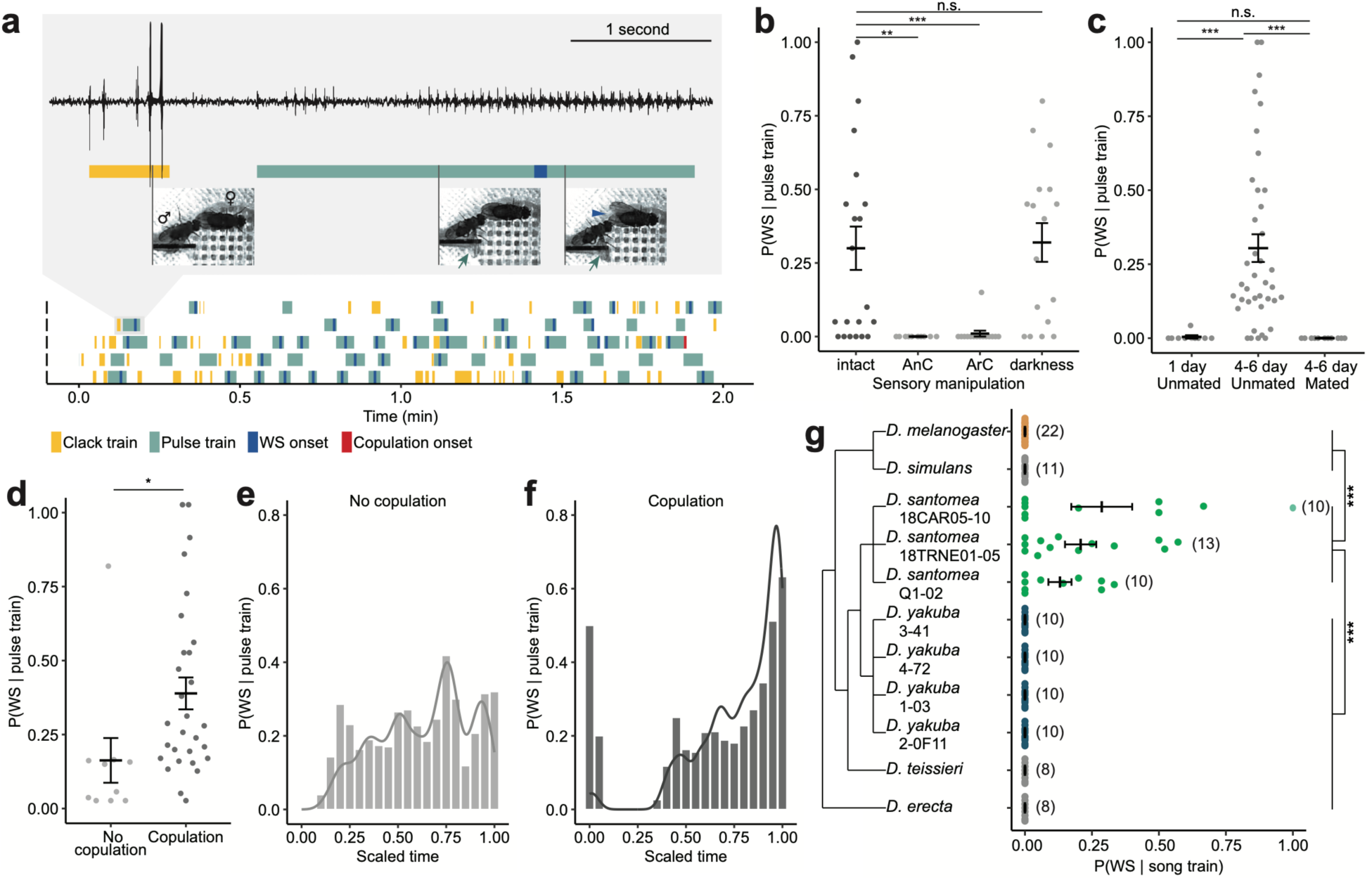
WS in *D. santomea* is a recently originated female receptive behavior in response to male pulse song. **a,** Representative behavioral ethograms of 2-min window in 5 courting *D. santomea* pairs. Gray box: zoom-in showing song trace, ethogram annotation, and still photos of a courting pair during a clack and a pulse train. Green arrows point to male single wing extension during a pulse train, and the blue arrowhead points to female WS. **b,** Probability of observing WS in response to a male pulse train in intact, antennae cut (AnC), and aristae cut (ArC) females, and in pairs recorded in darkness. **c,** Probability of observing WS in response to a male pulse train in females separated by age-related sexual maturity and mating status. 1 day old females are sexually immature. **d,** Probability of observing WS in response to a male pulse train in sexually mature (4-6 day old) ummated females, separated by whether the pair copulated during the recording period. **e,f,** Probability of observing WS in response to a male pulse train (bar, sliding windows of 0.1 width and 0.05 step size) over time and the corresponding density distributions (curve) in pairs that did not copulate (**e**) or copulated (**f**) during the recording period. Time was scaled for each pair such that 0.00 represents the start of recording, and 1.00 represents the end of recording (**e**) or the onset of copulation (**f**). **g,** Probability of observing WS in response to conspecific male courtship songs in the *melanogaster* subgroup. Sample sizes indicated in parentheses. Error bars show mean±SEM. Statistical significance tested with linear models. *** p<0.001, ** p<0.01, * p<0.05, n.s. not significant.

*Drosophila santomea* males produce two types of courtship songs: trains of louder clack generated by bilateral wing vibration, primarily during chasing, and trains of quieter pulses generated by unilateral wing vibration, often when females slow down to allow males to sing in close proximity^13,24–26^. We found that WS responded selectively to pulse and not clack trains (Fig. 1a). Consistent with the observation that female WS followed an auditory signal, removing a female’s antennae or aristae to abolish her hearing^27^ completely eliminated WS (Fig. 1b). In comparison, females invariantly performed WS in light *versus* dark conditions, showing that the production of WS does not depend on visual signals (Fig. 1b).

We further determined how WS is modulated by a female’s internal state of receptivity. In sexually mature unmated females, 30.4% of pulse trains elicited female WS. However, unreceptive females, either sexually immature or recently mated, rarely exhibited WS (Fig. 1c). Moreover, among the mature unmated females, those who had accepted a male’s copulation attempt responded with WS more frequently than those that did not, suggesting a correlation between WS probability and female receptivity to copulation (Fig. 1d). During the courtship interaction, a female continuously evaluates male quality based on his signals, which might influence her receptivity and inform her copulation decision. Indeed, we observed a major increase in WS probability leading up to copulation (Fig 1e,f), and 75.0% of the last pulse train before copulation elicited WS. Therefore, WS probability reflects not only female receptivity at the level of sexual maturity and mating status, but also temporal changes during the courtship interaction.

Given that WS behavior has not been previously reported in the *melanogaster* subgroup, we asked if WS represents a recent behavioral innovation in *D. santomea*. We therefore recorded receptive females from five species in this subgroup spanning ∼6 million years (Myr) of divergence^28^: *D. melanogaster*, *D. simulans*, *D. yakuba*, *D. teissieri*, and *D. erecta*. In none of these species did we detect WS (Fig. 1g; Spieth^19^ documented a 10° WS as an acceptance signal in female *D. simulans*, but we did not observe such behavior in our strain). We further sampled additional strains of *D. santomea* and its closest sibling species *D. yakuba*. Consistently, females from all *D. santomea* strains exhibited WS, while none from the *D. yakuba* strains did (Fig. 1g). This indicated that WS might be a fixed species difference instead of an intraspecific variation among *D. santomea* strains. *D. santomea* is endemic to the volcanic island of São Tomé, while *D. yakuba* is widely distributed in Africa^29^. We conclude that WS recently originated in the island species *D. santomea* when it diverged from *D. yakuba* about 0.4-1 Myr ago^30–32^.

### Function of WS as a receptive female signal

Female WS might be a social signal that actively modulates a male’s behavior or simply a facilitating act that exposes her genitalia and thereby assists a male’s licking and attempted copulation. To distinguish between these scenarios, we examined the effect of abolishing WS, by removing a female’s wings, on copulation success. A reduction in copulation success would be suggestive of WS’s signaling role, while an increase would be suggestive of WS’s facilitative role. We found that males paired with wing-cut females sang a similar amount of pulse trains (Extended Data Fig. 2a), but had a much lower copulation rate than those paired with intact females (Fig. 2a), suggesting that female WS is a functional signal. Consistent with WS being a species-specific signal, wing removal in *D. yakuba* and *D. melanogaster*, whose females do not perform WS, did not affect their copulation rate (Fig. 2a).

**Fig. 2:**
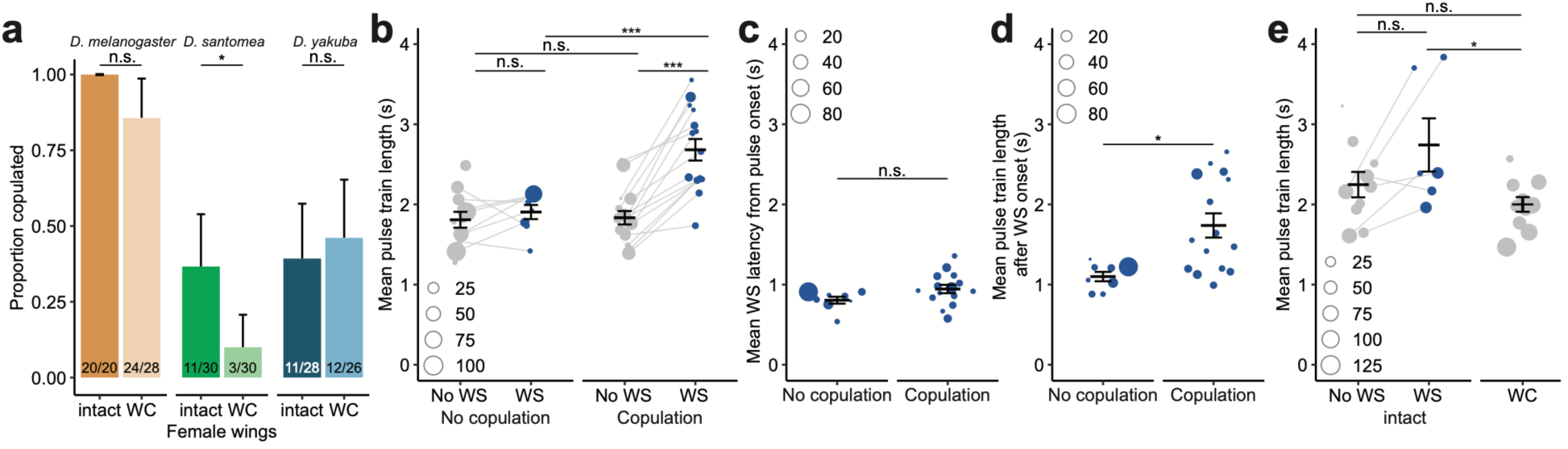
WS-dependent copulation success and song modulation in *D. santomea*. **a,** Proportion of pairs with intact or wing-cut (WC) females that succeeded in copulation in each species. Error bars represent the 95% confidence interval. Fractions at the base of each bar denote “number of pairs that copulated”/“total pairs tested”. Significance tested by Fisher’s exact test. **b,** Mean length of pulse trains separated by whether they elicited WS and whether the pair copulated during the recording period. Dot size corresponds to the number of pulse trains of each type in each pair. **c,d,** Mean latency of WS from pulse train onset (**c**) and mean pulse train length after WS onset (**d**), respectively, separated by whether the pair copulated during the recording period. In (**d**), the non-parametric Mann-Whitney U test is used to test for statistical difference between the two groups. **e,** Mean length of pulse trains in pairs with intact females, separated by whether they elicited WS, and in pairs with WC females. Only pairs that did not copulate during the recording period are shown. Dot size in **b-e** corresponds to the number of pulse trains of each type in each pair. Unless otherwise specified, error bars show mean±SEM and statistical significance was tested with linear models.. *** p<0.001, * 0.01<p<0.05, n.s. not significant.

We next sought to understand how WS, by communicating a female’s receptivity, alters a male’s behavior to influence the copulation outcome. We observed that WS coincided with a longer pulse train in pairs where females eventually accepted the males’ copulation attempts (Fig. 2b and Extended Data Fig. 2b), thus prompting two possibilities. Firstly, female WS motivates a male to sing longer pulse trains, with the male’s ability to adjust singing efforts predicting or directly affecting his copulation success. Alternatively, longer pulse trains are more potent at eliciting a female’s WS response, and males who produce these longer pulse trains have higher copulation success. We found that WS typically occurred shortly after the start of a pulse train, indicating that a female’s decision to display WS did not depend on hearing a long pulse train. Concordantly, the long pulse train associated with WS in copulated pairs resulted from continued singing after WS began (Fig. 2c,d and Extended Data Fig. 2c,d). In addition, we directly tested the impact of WS on the length of pulse trains by removing female wings to prevent WS. We found that the duration of pulse trains were comparable to those not associated with WS and significantly shorter than WS-associated pulses (Fig. 2e and Extended Data Fig. 2e). Taken together, WS serves as a functional female signal that promotes sustained pulse singing in males. The link between enhanced male singing efforts and copulation success further points to sexual selection favoring males who adeptly respond to the WS signal.

### Relationship between WS and VPO

To understand how the newly originated WS behavior is integrated into the pre-existing courtship ritual, we examined the relationship between WS and other female behaviors. Like WS, VPO (when a female extends her abdomen and pushes open her vaginal plates) was reported to be a response to male courtship song in receptive females in *D. melanogaster*^33^. Given the similarity between WS and VPO in both the external sensory stimulus and the associated female receptive state, we tested if the two behaviors are associated.

In this dataset, a pulse train could evoke WS and VPO simultaneously (VPO+WS; 39.5%), just VPO (VPO-only; 17.1%), or neither behavior (Neither; 43.4%). Thus, WS always co-occurred with VPO, and we never observed OE in sexually mature unmated females. Using SLEAP, a deep-learning based animal pose tracker^34^, we monitored changes in female abdomen length as a quantitative readout for VPO and wing angle for WS before, during, and after hearing a pulse train (Fig. 3a-c). The velocity and relative positions of the interacting sexes are shown in Extended Data Fig. 3. Mostly notably, the VPO+WS events revealed a linearly correlated increase (p<1×10^-10^, adjusted R^2^=0.980) in abdomen length and wing angle upon pulse song onset until the maximum abdomen length was reached (Fig. 3c). Nonetheless, many VPO events happened without WS. VPO+WS events showed significantly more intense VPO than VPO-only events, measured by the maximum extension of abdomen (Fig. 3d). The co-occurrence and quantitative scaling of WS with VPO, as well as its preferential association with more intense VPO, together suggest that WS is layered on top of the conserved behavior VPO to communicate non-identical social information, potentially a higher receptivity level, during the courtship interaction.

**Fig. 3:**
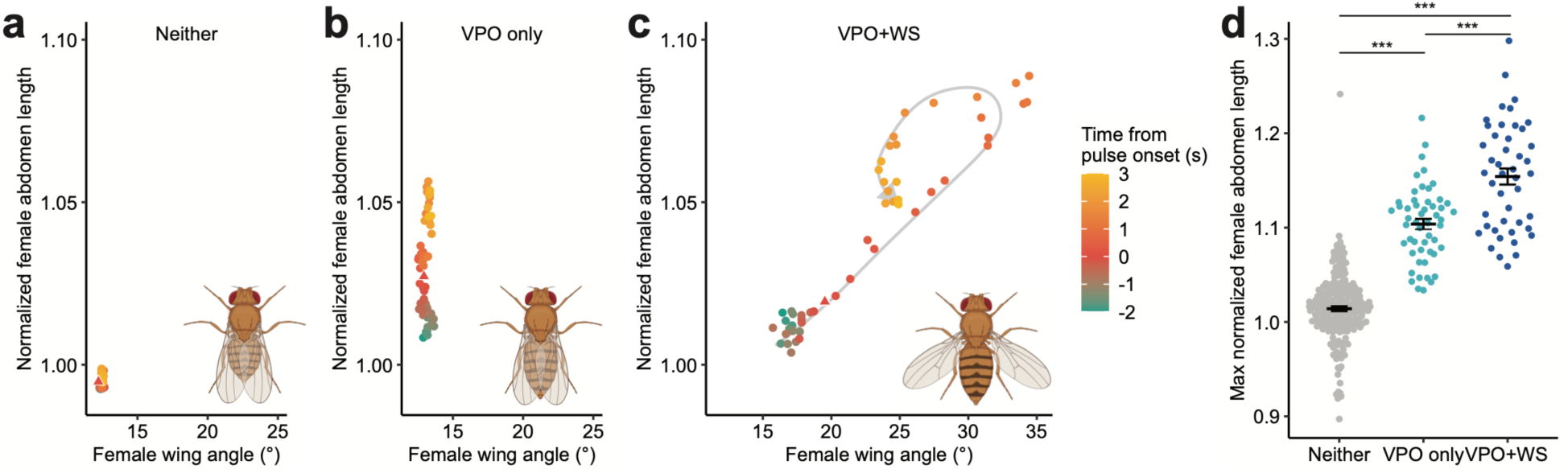
WS scales with VPO and co-occurs with VPO of higher intensity. **a-c,** Temporal relationship between normalized female abdomen length and female wing angle, averaged by event type: Neither (**a**), VPO only (**b**) and VPO+WS (**c**). Pulse onset is marked as a triangle. Gray arrow behind data points in (**c**) represents an approximate progression of data points. Inset diagrams illustrate each event type at maximum abdomen length and/or wing angle. **d,** Maximum normalized female abdomen length compared across all event types. Statistical significance was tested with linear mixed models using pair identity as a random effect. Error bars show mean±SEM. *** p<0.001.

### Co-option of VPO command neurons in WS

We hypothesized that WS emerged through modification of pre-existing female sexual circuits. Many circuit elements that encode female-typical behaviors express the sex determination gene *doublesex* (*dsx*), which undergoes splicing into sex-specific isoforms to guide the development of sexually dimorphic neural circuits^33,35–42^. In the brain of *D. melanogaster*, *dsx* neurons are organized in anatomically and functionally discrete neuronal clusters that function in various aspects of female reproductive behaviors^33,35,37–40,42–45^. For instance, pC1 neurons encode a female’s mating status^39,44,46^. Additionally, vpoDN (also known as pMN2) is a single pair of descending neurons that integrates the external and internal signals to function as a command control of VPO. They receive direct inputs from pC1 neurons and the male song-tuned auditory neurons in the brain, and project to the ventral nerve cord (VNC), primarily targeting the abdominal circuit^33^.

To compare the function of *dsx* brain neurons across species in relation to the origin of WS, we developed genetic tools that specifically labeled and manipulated *dsx* brain neurons in *D. santomea*, its sibling species *D. yakuba*, and the model species *D. melanogaster*. Specifically, we generated *dsx*-GAL4 alleles, using CRISPR/Cas9 genome editing, and brain-specific flippase transgenes^47^, which together restricted GAL4-dependent expression of effector genes to *dsx* neurons in the brain. The gross anatomy of *dsx* brain neurons labeled was similar across the three species (Fig. 4a). By expressing CsChrimson^48^, we optogenetically activated *dsx* brain neurons in isolated, freely-moving females and tracked their body coordinates using SLEAP^34^. Neural activation drove robust abdomen extension in all three species (Fig. 4b,c and Video 2). Based on findings in *D. melanogaster*, this abdomen phenotype can be readily explained by the activity of vpoDN in triggering VPO and/or the activities of DNp13 (also known as pMN1) and pC2l in triggering OE^33,40,42,43^. In contrast to the conserved abdomen phenotype, the same activation only triggered WS (manifested as an increased wing angle) robustly in *D. santomea* females (Fig. 4d,e and Video 2). Remarkably, decapitated females with only VNC neurites of vpoDN and DNp13 descending neurons activated (Fig. 4a) recapitulated both abdomen extension and wing angle results: females from all three species showed similar abdomen extension, but only *D. santomea* females displayed robust WS (Fig. 4f-i, Video 2). This result effectively restricted the neurons responsible for WS in *D. santomea* to the two candidate pairs. Unlike VPO and WS, OE represents a rejective female state^20,21,40,42^. Further, in natural behaviors of *D. santomea*, WS obligately co-occurs with VPO while never with the rejective behavior OE. Taken together, we inferred that activation of vpoDN elicited WS in *D. santomea*.

**Fig. 4:**
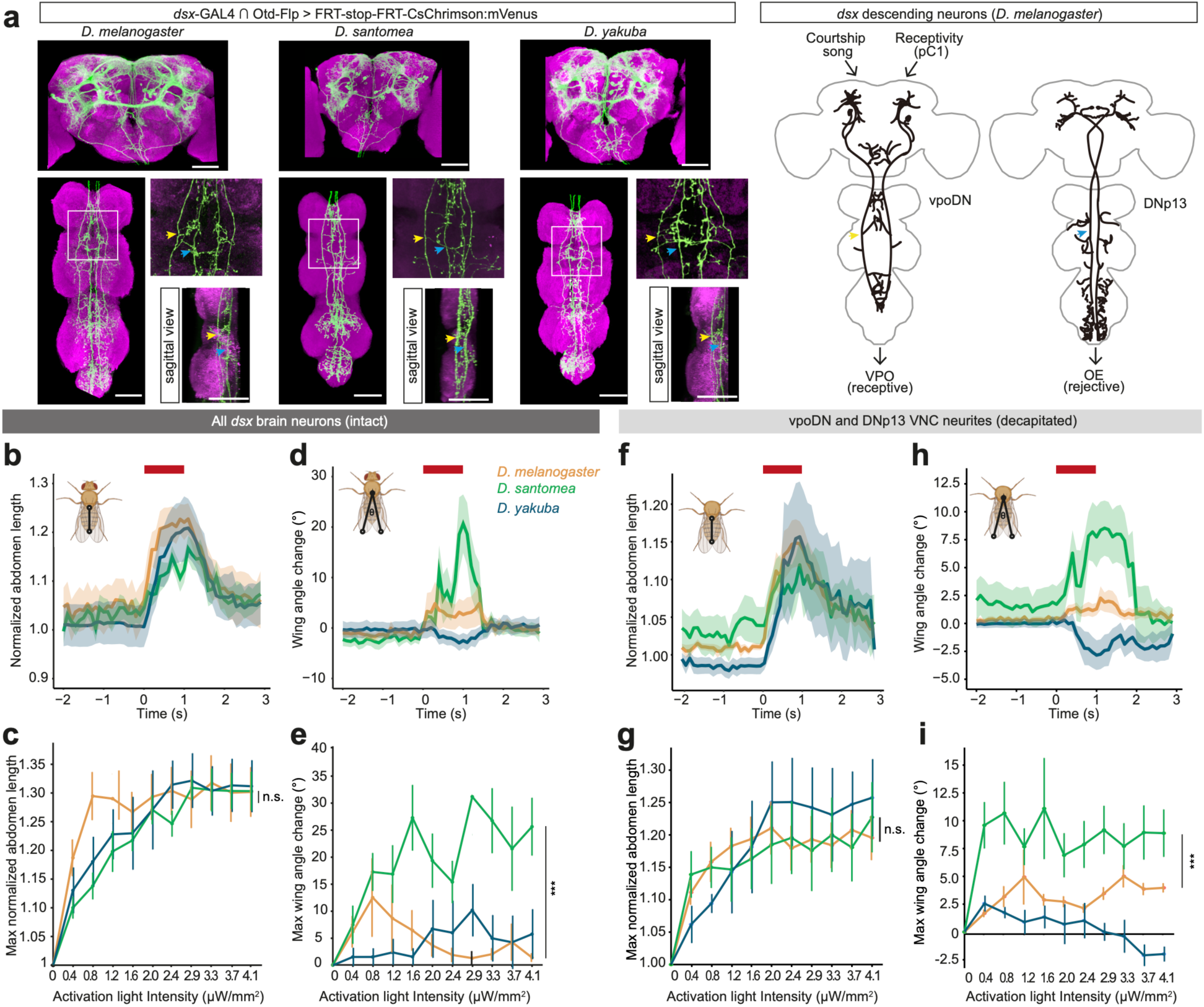
Activation phenotypes of brain *dsx* descending neurons across three species. **a,** Confocal images of female *dsx* brain neurons in the brain (top) and VNC (bottom) of each species. Green: mVenus; magenta: nc82. Only two pairs of neurons, vpoDN and DNp13^33,40^, project into VNC. Arrows highlight VNC projections of vpoDN (yellow) and DNp13 (blue). Scale bars: 50 µm. **b-i,** Behavioral phenotypes of optogenetically activating *dsx* brain neurons in intact (**b**-**e**) and decapitated (**f**-**i**) females of each species. **b,d,f,h:** Mean normalized abdominal length (**b**,**f**) and wing angle change (**d**,**h**) of intact females (**b**,**d**) at 1.6 µW/mm^2^ or decapitated females (**f**,**h**) at 0.8 µW/mm^2^. Activation window is denoted by red bars. Shaded areas represent the SEM. Inset diagrams illustrate how abdomen lengths or wing angles were measured. **c,e,g,i:** Maximum normalized abdomen length (**c**,**g**) and wing angle change (**e**,**i**) of intact females (**c**,**e**) or decapitated females (**g**,**i**) under each activation intensity. Wilcoxon rank sum tests were performed only between *D. melanogaster* and *D. santomea* (activation triggered female song in *D. yakuba*). Curve and error bars show mean±SEM. *** p<0.001, n.s. not significant.

Aside from the WS phenotype in *D. santomea*, we also observed behavioral changes in the other two species upon activating *dsx* brain neurons. In *D. yakuba*, females moved wings inward while generating a polycyclic song (Extended Data. Fig. 4a and Video 3), a behavior that has not been observed in wildtype *D. yakuba* in this study nor reported before. In *D. melanogaster*, there was a slight increase in the wing angle upon activation (Fig. 4d,h), contributed by a few females (30.0% intact, and 20.0% decapitated) exhibiting WS (Extended Data Fig. 4b and Video 4). Therefore, *D. melanogaster* has a latent circuit for WS.

### Latent potential of vpoDN to drive WS in *D. melanogaster*

Given the likely role of vpoDN in WS in *D. santomea*, we hypothesized that the activated WS phenotype in *D. melanogaster* also stemmed from the activity of vpoDN. Indeed, optogenetic activation of vpoDN neurons using a previously reported genetic reagent^33^ (Fig. 5a) induced VPO in all females and WS in 21.9% of females (Fig. 5b, Extended Data Fig. 5a,b, Video 5). We note that this vpoDN line has a different genetic background from the reagent labeling all *dsx* brain neurons. The idiosyncrasy of vpoDN in inducing WS across different genetic backgrounds of *D. melanogaster* suggested that it might be attributable to stochasticity during development. Environmental factors, such as a high temperature during development, can challenge the robustness of non-canalized developmental mechanisms and introduce stochasticity^49–51^. Hence, we tested the effect of developmental temperature, an impactful environmental factor on neuronal morphology and synaptic physiology^52–54^, on the efficacy of vpoDN activation in eliciting WS. Intriguingly, rearing flies at a high temperature of 29°C, relative to 23°C, during the larva and pupa stages drastically boosted vpoDN’s potential to elicit WS (Fig. 5c-g, Extended Data Fig. 5c-j) to 71.3% of females. This temperature effect was robustly manifested across different activation intensities (Fig. 5g and Extended Data Fig. 5c,d,g,h). In contrast, VPO was fully canalized regardless of the developmental temperature, and no major effect was observed for the proportion of responding females (100% *versus* 100%) or the extent of abdominal extension (Fig. 5d,f and Extended Data Fig. 5e,f). In sum, vpoDN has a latent potential to induce WS in *D. melanogaster,* and this potential is idiosyncratic and subjected to temperature-dependent developmental plasticity.

**Fig. 5:**
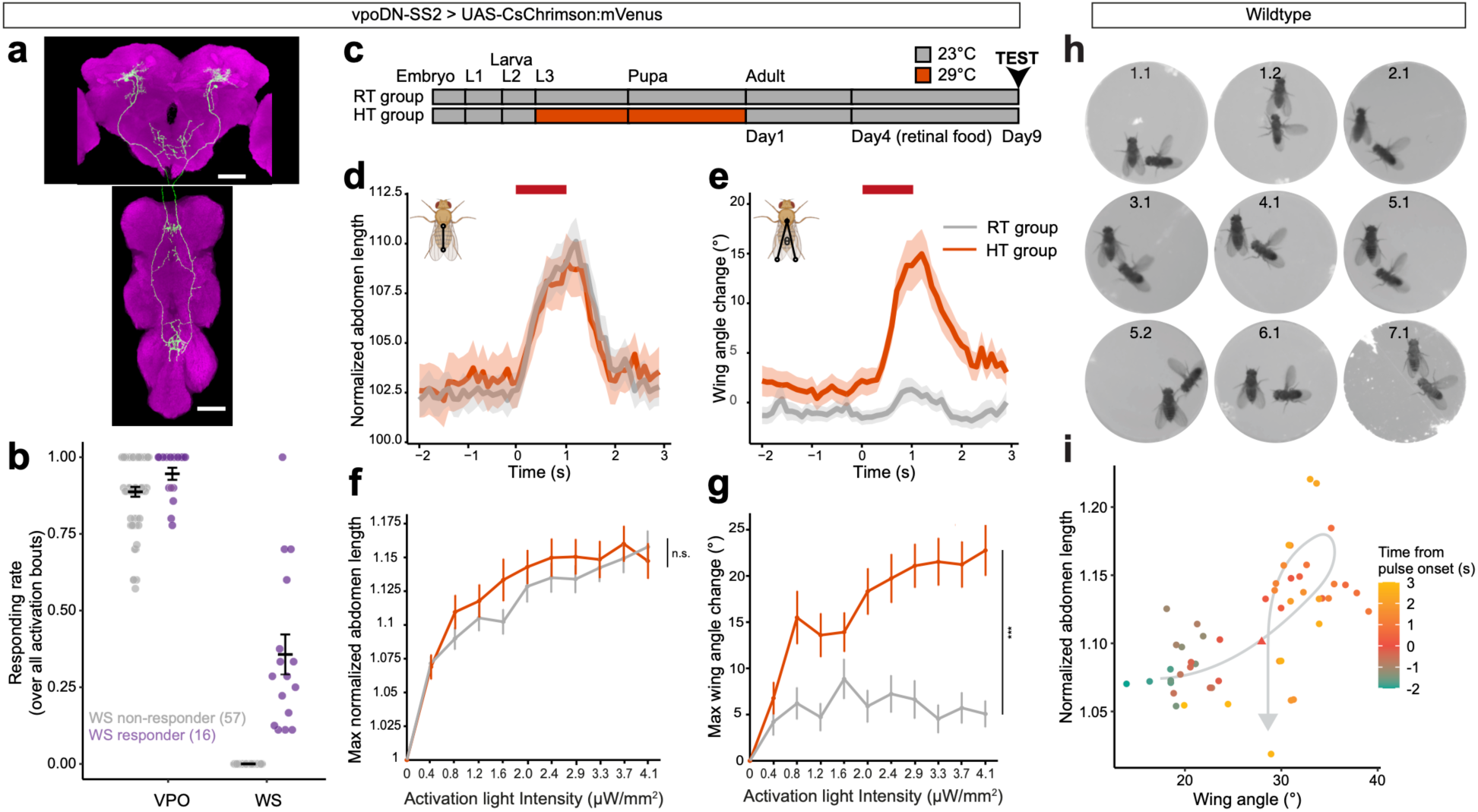
Idiosyncratic and plastic latent potential of WS in *D. melanogaster*. **a,** Confocal image of vpoDN neurons in *D. melanogaster* vpoDN-SS2 > UAS-CsChrimson:mVenus in female brain and VNC. Green: mVenus, magenta: nc82. Scale bars: 50 µm. **b,** Proportion of VPO and WS events in response to 10 activation bouts with intensities ramping from 0.4 to 4.1 µW/mm^2^. Each dot represents an individual. Color represents whether an individual was scored as a WS responder (purple) or not (gray). Error bars show mean±SEM. **c,** Schematic of how room temperature (RT) and high temperature (HT) groups were generated. **d,e,** Mean normalized abdomen length (**d**) and wing angle change (**e**) of HT and RT flies at 4.1 µW/mm^2^. Activation window is denoted by red bars. Shaded areas represent the SEM. Inset diagrams illustrate how abdomen lengths or wing angles were measured. **f,g,** Maximum normalized abdomen length (**f**) and wing angle change (**g**) under each activation intensity. Curve and error bars show mean±SEM. Wilcoxon rank sum tests were performed between RT and HT across all activation intensities. *** p<0.001, n.s. not significant. **h,** WS onset frame of each of the 9 WS events observed in 7 courting wildtype pairs. Numbers denote “pair ID”.“event ID”. **i,** Temporal relationship between normalized abdomen length and wing angle, averaged across all WS events. Pulse onset is marked as a triangle. Gray arrow behind data points represents an approximate progression of data points.

Given the latent potential of vpoDN to elicit WS, we examined whether wildtype *D. melanogaster* females occasionally exhibited WS in a way that was overlooked in previous studies. In total, we identified 9 WS events contributed by 7 females from assaying the courtship interactions of 141 pairs (7 of 105 pairs with females raised at 29°C, and 0 of 36 pairs with females raised at 23°C, Fig. 5h, Video 6). All WS events co-occurred with VPO, and 4 out of 9 immediately preceded copulation. Also mirroring the natural WS behavior in *D. santomea*, there was a positive temporal correlation between female wing angle and abdominal length upon the onset of male singing (Fig. 5i). Thus, wildtype *D. melanogaster* females perform WS at very low frequency in certain conditions.

### *Drosophila santomea* WS is a recurrent variant of a receptive female behavior

Beyond the *melanogaster* subgroup, female WS has been reported in a few other species within the *Sophophora* subgenus, and more broadly in the *Drosophila* subgenus as a pre-copulatory acceptance signal that initiates copulation^19,22,23^. Whether females also perform WS during the courtship interaction as in *D. santomea*, and how WS is associated with VPO, have not been explicitly investigated in this broader phylogenetic context. Therefore, we surveyed 22 species in the *Sophophora* and *Drosophila* subgenera for VPO and WS during courtship and before copulation (Fig. 6, Video 7). As expected, VPO was a conserved female behavior observed in all species. In contrast, WS was sparsely represented in the *Sophophora* subgenus, with repeated turnovers in multiple lineages. In species with WS, these events were not specifically linked to copulation: in the *Sophophora* subgenus, WS was more commonly seen during the courtship interaction than right before copulation; whereas in the *Drosophila* subgenus, WS appeared obligatory prior to copulation but was also observed during courtship. Therefore, WS can be broadly characterized as a receptive signal that communicates females’ sexual interests instead of an acceptance signal that green-lights copulation, while the precise social context and receptivity state that WS represents may vary across species. Additionally, both the association and the decoupling between WS and VPO are common behavioral features across the phylogeny (Fig. 6), suggesting the possibility of shared circuit mechanisms that facilitate the recurrent evolution of WS.

**Fig. 6:**
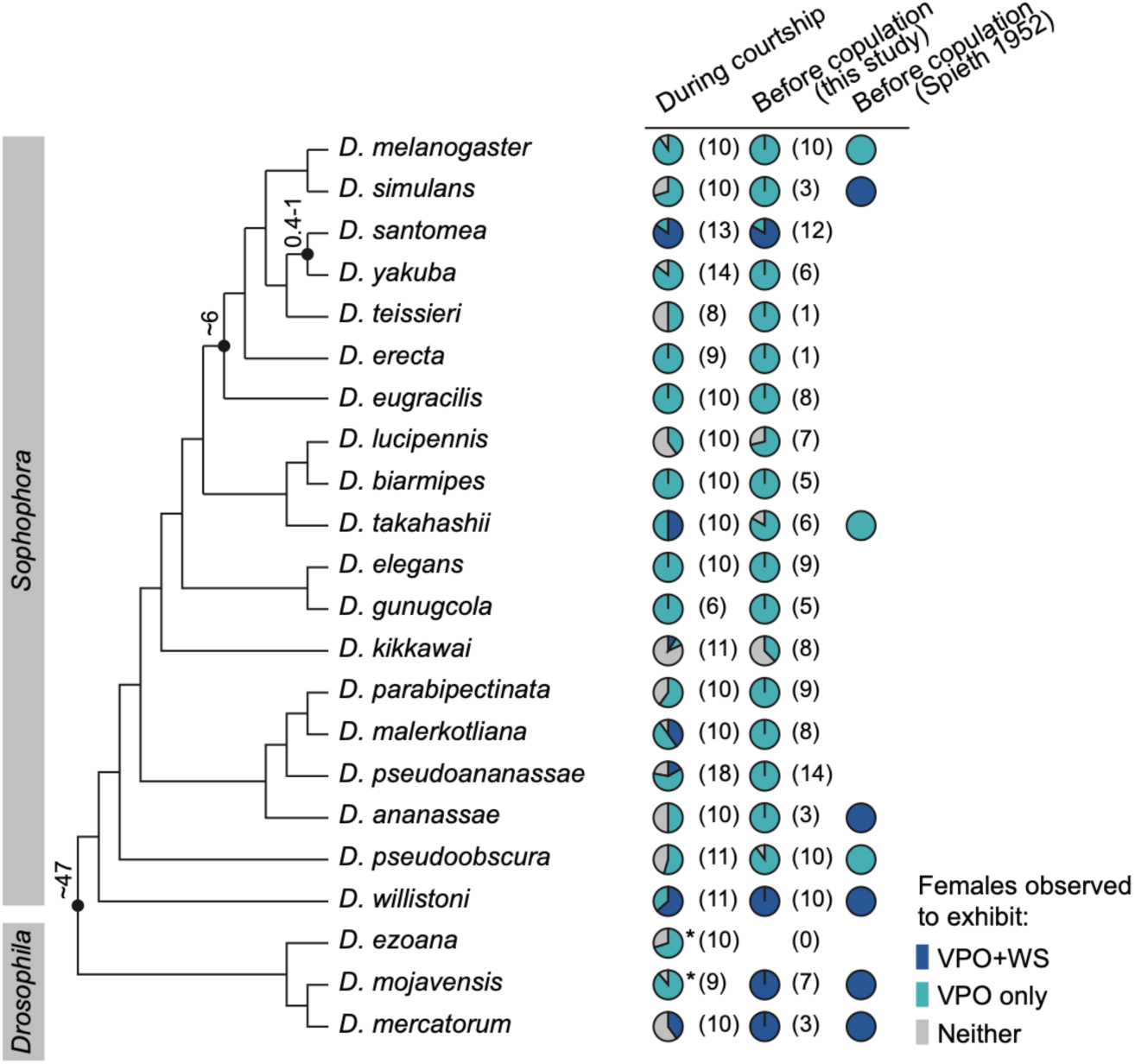
WS is found in multiple *Drosophila* lineages. First two columns: proportion of females observed to exhibit VPO+WS, VPO, or neither behavior in pairwise matings under the designated social context (above). Asterisk denotes species where females were observed to sing courtship duets with males, and song-independent WS behavior was not observed. Sample sizes are indicated in parentheses. Last column: published results from Spieth 1952 on whether the pre-copulatory acceptance behavior is VPO or VPO+WS. Numbers next to key nodes indicate estimated divergence times in Myr^28,30–32,86^. Phylogeny is based on ^29,86–90^.

## Discussion

Historically, studies on the evolution of mating behaviors have predominantly focused on male signals. An emerging perspective shift repositions females as active participants in the dynamic courtship interaction and not passive receivers of male signals^4,7,55,56^. Nonetheless, female courtship behaviors remain much under-characterized, and little is known about how they originate or evolve. Here, we show that *D. santomea* WS, a species-specific female behavior originated within the last 0.4-1 Myr, is layered on top of conserved elements of the dynamic courtship interaction to affect male behaviors and direct mate selection. WS *in D. santomea* serves both as a female response to a male’s cue (the pulse song) and as a signal of her sexual interests, thereby influencing the male’s subsequent actions. Intriguingly, whether a male is capable of increasing his efforts accordingly is a predictor of his chances of copulation, supporting a pivotal role of female sexual behavior in organizing a social feedback loop upon which sexual selection operates. *Drosophila santomea* co-localizes with the sibling species *D. yakuba* in a hybrid zone on the island of São Tomé^29,57,58^. As such, WS might be a key phenotype in the reproductive isolation in these two naturally hybridizing species.

### Expression of a new behavior in the appropriate social context

Capitalizing on this recently originated female behavior, our neural circuit comparisons across species shed light on the neural mechanisms by which a new social behavior may originate. The co-option of the VPO command neurons vpoDN, which integrate both sensory and motivational information^33^, would allow a receptive female hearing a potent male song to express the new behavior WS, and thereby communicate her interests to the male. Descending neurons like vpoDN act as a critical information bottleneck that compresses high-dimensional brain dynamics to low-dimensional commands that interface with motor circuits^59,60^. The co-option of vpoDN in WS suggests that existing descending pathways might be restrictive neural substrates favored by evolution to drive new behaviors, because they readily permit the expression of newly originated behavior in a meaningful social context.

As vpoDN evolved from a uni-functional node that only drives VPO to a possibly bi-functional one that drives both VPO and WS, we further consider how WS can be encoded in a way that communicates non-identical social information from the ancestral VPO signal. Social behaviors, such as mating and aggression, may involve a combination of behaviors that are associated with graded states of drive^61–63^. We showed that WS tended to co-occur with more intense VPO in wildtype *D. santomea*, and the intensity of VPO increased with the activation intensity of vpoDN in *D. melanogaster*. Therefore, compared with VPO, the expression of WS might involve a higher level of vpoDN activity. Because vpoDN activity reflects female receptivity by receiving excitatory inputs from pC1 neurons^33^, it is possible that WS is differentially gated from VPO by vpoDN activity to represent a higher receptivity level. Alternatively, modulatory inputs independent of vpoDN could contribute to the differential expression of WS and VPO in natural behaviors. Future testing of the hypotheses would benefit from genetic tools that specifically label vpoDN in *D. santomea*.

### Latent circuit potential facilitates the evolution of new behaviors

How does a socially informed behavioral decision lead to a new motor action? The co-option of vpoDN to elicit WS in *D. santomea* suggests that it must be functionally coupled with a wing motor circuit. When the VNC neurites of vpoDN and DNp13 were activated in decapitated flies (who lacked inputs from the central brain), WS was elicited most robustly in *D. santomea*. Therefore, the functional connection between vpoDN and the downstream WS motor circuit has evolved to be more potent in *D. santomea*, pinpointing the neural substrates behavioral divergence. Meanwhile, in *D. melanogaster*, vpoDN activation sometimes induced WS, and wildtype females displayed WS on rare occasions, suggesting that this connection is not a *de novo* but rather an ancestral feature that remains largely latent yet potent. Previous studies reported that sex- and developmental stage-typical behaviors can be experimentally induced, suggesting that latent potentials may broadly exist in the nervous system^64–69^, serving as raw substrates that fuel the rapid evolution of new behaviors. If so, we anticipate that species-specific behaviors may commonly exist in closely-related outgroup species in primitive prototypes that are occasionally expressed under certain conditions.

The absence of WS within the *melanogaster* subgroup (besides *D. santomea*) and its presence in some species outside of the *melanogaster* subgroup raises the possibility that WS in *D. santomea* evolved from the reactivation of a vestigial circuit that has lost the WS function at least ∼6 Myr ago^28^. Indeed, neurons for lost behaviors, such as wing motoneurons in flightless grasshoppers, can survive millions of years’ evolution^66,70^. However, it is also possible that the ancestral connection between vpoDN and the wing motor circuit is not a vestigial feature on the way of degeneration but rather actively maintained to serve an as-yet undefined function. In all three species we examined, vpoDN projects to the mesothoracic neuromere, where they branch dorsally and medially to innervate the tectum, potentially permitting a contact with the wing circuit. One hypothesis is that this connection is required for movement coordination, such as the engagement of wing muscle to sustain a proper posture during VPO. In this scenario, the circuit configuration, maintained by selective pressures unrelated to WS, potentiates the recurrent evolution of WS within *Sophophora*.

### Species difference, idiosyncrasy, and plasticity in behaviors

In *D. melanogaster*, vpoDN’s ability to drive VPO constitutively *versus* WS as a latent potential presents a comparison. The former is fully penetrant and canalized: all individuals displayed VPO in response to vpoDN activation, and the response was unaffected by developmental temperature. In contrast, the latter is idiosyncratic and plastic: only some individuals responded with WS, and the response was strongly influenced by developmental temperature. Notably, here, the species difference, idiosyncrasy, and plasticity in behaviors, despite operating at different levels, all reflect phenotypic variations of the same neural circuit substrates, highlighting the lability of ancestral circuits in encoding new behavioral prototypes. Such a labile circuit can then be refined to encode stably expressed behaviors when a selective pressure is present. With a prototypic circuit in place, minor modifications on the weights of local excitations or inhibitions might be sufficient to allow vpoDN to stably engage the wing motor circuit. Extensive resources in the model species *D. melanogaster*, such as EM connectomes and neurogenetic tools^71–75^, will facilitate future characterizations of the organization and evolution of the underlying circuits.

By developing a comparative behavioral and neural paradigm to investigate the origin of new behaviors, our results revealed how the ancestral nervous system potentiates such changes and shapes the trajectories of behavioral evolution. The themes emerging from this study, such as co-option, ancestral potential, and the plasticity of prototypic phenotypes, converge with Evo-Devo concepts that focus on morphological evolution^76–79^. For example, analogous to the origin of WS, the recurrent evolution of ‘supersoldiers’ in the ant Genus *Pheidole* occurred via the actualization of an ancestral developmental potential, where large supersoldier-like anomalies are occasionally found in nature and can be artificially induced by hormonal manipulation in species lacking a supersoldier caste^80^. The dissection of neural mechanisms underlying the origin of new behaviors contributes to the synthesis of principles unique for behavioral evolution as well as a unifying conceptual framework for phenotypic evolution^81^.

## Material and Methods

### Fly stocks

Flies were maintained on cornmeal-agar-yeast medium (Fly Food B, Bloomington Recipe, Lab Express) at 23°C and 50% humidity on a 12 hour light/dark cycle, unless otherwise specified. All fly stocks used in this study are listed in Supplementary Table 1.

### Generation of transgenic flies

The *dsx*-GAL4 knock-in alleles in *D. santomea*, *D. yakuba*, and *D. melanogaster* were generated using an identical design of CRISPR/Cas9 genome editing to replace the majority of the first coding exon (second exon) of *dsx* in-frame with T2A-GAL4 followed by a Mhc-DsRed marker. The donor plasmids were constructed by concatenating a left homology arm, T2A, GAL4 (amplified from pBPGUw, addgene #17575^82^), a Mhc-DsRed marker^83^, a right homology arm, and a 1.8 kb backbone using Gibson Assembly (NEB). A pair of guide RNAs (gRNAs) were cloned into the pCFD4 vector. A cocktail containing the donor plasmid (200 ng/ul), pCFD4 plasmid (200 ng/ul), and *in vitro* transcribed Cas9-nanos 3’UTR mRNA (200 ng/ul) was injected into fly embryos. The Otd-Flp lines, which drive Flp expression exclusively in brain, were generated by inserting the pBpGuW-Otd-nls:FLPo plasmid^47^ into the 2253 attP landing site on the third chromosome in *D. santomea* and the 2285 landing site on the third chromosome in *D. yakuba*^84^ using attB/P φc31 integrase system. The FRT-stop-FRT-CsChrimson:mVenus lines were similarly generated by inserting the pJFRC300-20XUAS-FRT>-dSTOP-FRT>-CsChrimson-mVenus plasmid^85^ into the 2253 site in *D. santomea* and the 2180 landing site on the second chromosome in *D. yakuba*^84^. The *dsx*-expressing brain neurons were labeled and activated using a genetic intersection of *dsx*-GAL4 and Otd-Flp to drive the expression of CsChrimson:mVenus, where the brain-specific recombinase (Otd-Flp) excises a transcriptional stop cassette (FRT-stop-FRT) to enable the transcriptional control of UAS-CsChrimson:mVenus under *dsx*-GAL4 only in brain. All injections were performed at Rainbow Transgenic Flies using a standard protocol.

### Preparation of flies for behavioral assays

Flies in the *Sophophora* subgenus used in behavioral assays were collected within a few hours of eclosion and kept in single-sex vials with 10-15 flies in each vial. Males were separated into individual vials at least 3 days before recording. All flies were 3-6 days old at the time of the assay, with the exception of the 1 day old sexually immature females in Fig. 1c. Mated females in Fig. 1c were generated by mating each female with a wildtype male 24 hr prior to recording. Flies in the *Drosophila* subgenus were collected the same way, but allowed age for 10-12 days before recording, and males were separated into individual vials at least 8 days before recording.

In optogenetic activation experiments, females were collected the same way as wildtype females but kept on medium supplemented with 0.2 mM all trans-retinal (Sigma Aldrich) in the dark for 5 days until recording. In wing-cut, antennae-cut, or aristae-cut experiments, female wings, antennae, or aristae, respectively, were removed bilaterally under CO_2_ anesthesia using micro scissors 3 days before recording. Control females were also subjected to CO_2_ anesthesia alongside the experimental females. Each female was exposed to CO_2_ for less than 3 minutes. In decapitation experiments, females were cut at the neck using micro scissors under CO_2_ anesthesia 30 minutes before the recording, and were allowed to recover in a vial with food until the recording. In temperature manipulation experiments, *D. melanogaster* females were either grown according to the presented scheme (Fig. 5d) or at 29°C throughout development (Fig. 5i,j).

### Behavioral recording

Two cameras (FLIR BFS-U3-200S6M-C, Edmud optics #11-521) with 50mm lens (Edmud optics #63-248) were used to record videos at 10 Hz. For audio recording (Figs. 1,2,6), we used a 3D printed behavioral chamber with beveled circular arenas fitted with fine mesh below. The arenas measured 10 mm in diameter and 3 mm in height. *Drosophila ezoana* pairs (Fig. 6) were placed in arenas measuring 15 mm in diameter to accommodate their larger body size. Each arena was placed on top of a microphone in a custom 96-channel recording apparatus, SongTorrent, that enabled simultaneous audio (5 kHz) and video recording. To optimize video recording (Figs. 3-5) for behavioral tracking, we used acrylic behavioral chambers with circular arenas that measured 10 mm in diameter and 3 mm in height and did not perform audio recording. In all recordings of female-male pairs, the flies were separated by a divider until the start of the recording. Wildtype flies were recorded for 20 min. Optogenetic flies were recorded for the duration of the activation scheme.

In optogenetic activation experiments, flies were allowed to see in blue light and recorded under infrared light (850 nm). Red light (635 nm) was used for activation following a programmed cycle. An activation cycle consisted of 10 activation bouts with increasing intensity, and each 1 s bout was interspersed with 9 s intervals. The only exception to this activation scheme was the *D. yakuba* audio recording (Extended Data Figure 4a), which had 10 s activation bouts and 10 s intervals. Activation intensity gradient (in µW/mm^2^) was as follows: 0.4, 0.8, 1.2, 1.6, 2.0, 2.4, 2.9, 3.3, 3.7, and 4.1.

### Behavioral tracking

SLEAP (v1.2.0a6^34^) was used to track the behavior of interacting pairs (in wildtype experiments) or individual females (in optogenetic activation experiments). For pairs, we tracked the head, thorax, abdomen, and each of the wing tips for each fly. A classifier was trained using the multi-animal top-down mode with the default settings and the following modifications: anchor part=thorax, rotation min and max angles=-180 and 180, scale=TRUE, contrast=TRUE. Inference was run using a simple tracker with default settings and the following modifications: 2 instances/frame, cull to target instance count=TRUE, all nodes are used for tracking, and connect single track breaks=TRUE. The onset of pulse trains were used as key frames, and we focused on the interval between 2 s before to 3 s after the pulse onset. Manual adjustments were made wherever necessary.

For individuals, we tracked the head, thorax, abdomen, tip of the external genitalia, and each of the wing tips. A single classifier was trained using the single animal model with default settings, and intact and decapitated *D. melanogaster, D. yakuba and D. santomea* were included in the training dataset. Inference was run using a simple tracker with 1 instance/frame. We focused on 2 s before and after each 1 s activation bout. Manual adjustments were made wherever necessary.

Tracking data was exported as HDF5 files and analyzed in Python (v3.8.13) and R (v4.2.2) to calculate parameters such as the female wing angle and abdomen length. In wildtype recordings, female abdomen length was normalized to the baseline of each female, calculated as the mean abdomen length across the 20 frames (2 s) before each pulse onset. In optogenetic recordings, each female’s baseline abdomen length used for normalization was calculated as the mean abdomen length over the 100 frames (10 s) before the first activation bout. Wing angle change was calculated by subtracting the observed wing angle by each female’s baseline wing angle, which was calculated as the mean wing angle over the 100 frames before the first activation bout.

### Behavioral analysis

#### Probability of WS in response to male song

The custom Matlab software Tempo (https://github.com/JaneliaSciComp/tempo) was used to annotate male songs, and when applicable, female WS in response to song. In Figs. 1c-f, 2b-e, and Extended Data Fig. 1 and 2, all pulse trains and WS were annotated manually. In Figs. 1b,g, if a male produced 20 or fewer pulse trains (or song trains in *D. melanogaster* and *D. simulans*), all pulse/song trains were annotated; otherwise 20 pulse/song trains were randomly sampled. Each pulse/song train’s co-occurrence with WS was then recorded.

#### Wildtype behavior

In wildtype recordings of non-*D. santomea* species (Fig. 5h,i and Fig. 6), full recordings were carefully examined for VPO and WS. To qualify as WS, a putative female wing extension behavior must occur in response a male courtship song and co-occur with VPO. These criteria were imposed to disambiguate WS from female wing flicking, grooming, and balancing after jumping. If VPO was not observed, it was typically associated with limited courtship history, non-ideal positioning of the female, and scoring challenge due to the subtlety of VPO in certain species.

#### Activation experiments

For vpoDN activation experiments in *D. melanogaster*, WS behaviors were manually identified by detecting wing angle changes in response to activation bouts. Responders (Fig. 5b and Extended Data Fig. 5c,d) were defined as females with at least one confirmed WS out of 10 activation bouts in an activation cycle. Responses to activation such as grooming, jumping and turning, and responses with low SLEAP tracking quality were considered invalid. Individuals with more than 5 invalid responses in the activation cycle were removed from Fig. 5b and Extended Data Fig. 5c,d. In Fig. 5b, invalid events were excluded when calculating WS and VPO rates of each individual. Notably, both intact and decapitated *D. santomea* females exhibited leaning and flipping over more frequently than the other species upon activation, possibly suggesting their lower resistance to activation. When females were not standing still, especially when leaning, SLEAP tended to underestimate the wing angle.

### Immunostaining

Female brains and VNCs were dissected in 1X Phosphate Buffered Saline (PBS; Thermo Fisher) within 50 minutes of ice anesthesia, fixed with 4% paraformaldehyde (PFA) for 35 min at room temperature rinsed 3 times in PBS with 1% Triton X-100 (PBTX), then blocked with 5% normal goat serum (NGS) in PBTX for 1.5 hours. Samples were incubated in primary antibodies (diluted in 5% NGS) at 4°C overnight. Samples were then washed with PBTX 3 times for 30 min each, and incubated with secondary antibodies (diluted in 5% NGS) at 4°C overnight. After 3 washes with PBTX, each 30 min, the samples were mounted with ProLong^TM^ Gold antifade reagents (Fisher Scientific; Cat.#: P36931) on poly-L-lysine coated coverslips, and sealed on all slides with nail polish. Primary antibodies used were: chicken-anti-GFP (1:600, ab13970, Abcam), mouse-anti-nc82 (1:30, DHSB). Secondary antibodies used were: goat-anti-chicken/ AF488 (1:500, A-11039, Thermo Fisher), goat-anti-mouse/AF568 (1:500, A-11031, Thermo Fisher). Confocal images were taken on a Leica DMi8 microscope with a TCS SP8 Confocal system at 40x, and processed with VVDViewer (v1.6.4).

### Selection of vpoDN split-GAL4 lines

Three vpoDN split-GAL4 lines (SS1, SS2, and SS3)^33^ were each crossed to UAS-CsChrimson:mVenus flies to assess their ability in eliciting VPO upon optogenetic activation. Behavioral recording, tracking, and analysis were performed as described above. The line vpoDN-SS2 was chosen for further experiments as it had the most robust abdomen extension phenotype (Extended Data Fig.5a).

### Statistical Analysis

Data analysis was performed in Python (v3.8.13) with the following packages: h5py (v3.6.0), numpy (v1.23.5), scipy (v1.8.0), and pandas (v1.4.2), and R (v4.2.2) with the following packages: tidyverse (v1.3.2), lme4 (v1.1-31), emmeans (v1.8.3), and lmerTest (v3.1-3). Scripts are available by request. Linear models and linear mixed models (to account for replicate effects and repeated measurements from the same subjects) were used to determine statistical significance. Variables that were proportions were arcsine-square root transformed to stabilize the variance. When there was significant deviation from the assumptions of linear models, non-parametric alternatives were used. Where relevant, p-values were adjusted for multiple testing using the Bonferroni correction.

## Acknowledgements

We thank Marc Schmdit, Troy Shirangi, Jan Clemens, and Ding lab members for discussion and comments on the draft. We thank David Stern lab for sharing genetic reagents, and David Anderson lab and Gerry Rubin lab for sharing plasmids for generating genetic reagents. We thank Steve Sawtelle for help with the behavioral recording system. This project was supported by Searle Scholarship and NIH grant R35GM142678 to Y.D. *Drosophila* illustrations were created with BioRender.

## Author contributions

D.S.C, M.L., and Y.D. conceived the study. D.S.C and M.L. collected and analyzed behavioral data. I.P.J. and M.L. collected imaging data. Y.D. and M.L. generated genetic reagents. F.S., G.H.C., and A.J.B. assisted annotation of behavioral data. D.R.M. and A.A.C. contributed fly strains and provided inputs to the manuscript. D.S.C, M.L., and Y.D. wrote the paper.

## Extended data

**Extended Data Fig. 1: Full-length behavioral ethograms of *D. santomea* courting pairs, including the ones shown in** **Fig.1a**.

Each row corresponds to one courting pair, with the numbers to the left representing their pair ID.

**Extended Data Fig. 2: Additional behavioral analyses linked to** **Fig. 2** **showing WS-dependent modulation of pulse train length.**

**a,** Number of pulse trains per minute that males produced when paired with intact or wing-cut (WC) females. Only data from pairs that did not copulate during the recording period are shown.

**b,** Length of pulse trains separated by whether they elicited WS and whether the pair copulated during the recording period.

**c,d,** Latency of WS from pulse train onset (**c**) and the length of pulse train after WS onset (**d**), respectively, separated by whether the pair copulated during the recording period. In (**d**), the non-parametric Mann-Whitney U test is used to test for statistical difference between the two groups.

**e,** Length of pulse trains in pairs with intact females, separated by whether they elicited WS, and in pairs with WC females. Only data from pairs that did not copulate during the recording period are shown.

Unless otherwise specified, error bars show mean±SEM, and statistical significance was tested with linear mixed models using pair identity as a random effect. *** p<0.001, ** p<0.01, * p<0.05,

n.s. not significant.

**Extended Data Fig. 3: Female and male behavioral parameters during pulse events.**

**a,** Mean male extended wing angle, male velocity, female velocity, normalized female abdomen length, female wing angle, and distance between male and female thoraces, separated by event type. Pulse onset is marked with a vertical gray line. Shaded areas represent the SEM.

**b,** Male position in female-centered coordinates during all pulse events, separated by event type. The female is represented by a triangle in the center, with the head pointing up. Each ring represents 1 mm. Male position at pulse onset is marked as a red triangle.

**c,** Distance between male and female thoraces at pulse onset compared across all event types. **d,** Male angle relative to the female body axis compared across all event types. At 180°, the male is directly behind the female.

Error bars show mean±SEM.

**Extended Data Fig. 4: Additional behavioral phenotypes of activating *dsx* neurons in *D. yakuba* and *D. melanogaster*.**

**a,** Audio traces upon activating *dsx* brain neurons in decapitated *D. yakuba* females using a 1s (left) or a 10s (right) activation scheme with a light intensity of of 4 µW/mm^2^. Each row represents one individual. Three examples of female song events (right) show the production of polycyclic signals with stereotypic waveforms in variable lengths.

**b,** Wing angle change in intact *D. melanogaster* females at 1.6 µW/mm^2^, and decapitated females at 0.8 µW/mm^2^. Activation window is denoted by red bars. Each line with a different color represents one individual. Individuals showing WS response in this activation window are highlighted with thicker lines.

**Extended Data Fig. 5: Additional behavioral analyses associated with** **Fig. 5** **showing developmental temperature-dependent modulation of WS phenotype upon vpoDN activation.**

**a,b,** Mean normalized abdominal length (**a**) and wing angle change (**b**) of *D. melanogaster* vpoDN-SS1, vpoDN-SS2 or vpoDN-SS3 > UAS-CsChrimson:mVenus females at 4.1 µW/mm^2^. Activation window is denoted by red bars. Shaded areas represent the SEM. Inset diagrams illustrate how abdomen lengths or wing angles were measured.

**c,d,** Wing angle changes of *D. melanogaster* vpoDN-SS2 > UAS-CsChrimson:mVenus females from RT group (**c**) and HT group (**d**) across 10 activation bouts with intensity from 0.4 to 4.1 µW/ mm2. Each row represents one individual. Each column represents a 5-second period centered at 1-second activation window (red bar) with light intensity listed above. Frames with low tracking quality are shown in gray. The manually scored responders are noted on the right side. **e-h,** Mean normalized abdominal lengths (**e**,**f**) and wing angles (**g**,**h**) of *D. melanogaster* vpoDN-SS2 > UAS-CsChrimson:mVenus females from RT group (**e**,**g**) and HT group (**f,h**) at different activation intensities denoted by colors.

**i,j,** Wing angle changes of *D. melanogaster* vpoDN-SS2 > UAS-CsChrimson:mVenus females from RT group (**i**) and HT group (**j**) at 3.3 µW/mm^2^. Ten individuals were randomly selected in the plot and each line represents one individual.

## Supplementary information

Video 1: Exemplar events of WS in wildtype *D. santomea* pairs.

Video 2: Typical behavioral phenotypes of activating *dsx* brain neurons in intact and decapitated *D. santomea*, *D. melanogaster*, and *D. yakuba*.

Video 3: Female singing induced by activating *dsx* brain neurons in *D. yakuba*.

Video 4: Occasional WS induced by activating *dsx* brain neurons in *D. melanogaster*.

Video 5: Occasional WS induced by activating vpoDN in *D. melanogaster*.

Video 6: Exemplar events of WS in wildtype *D. melanogaster* pairs.

Video 7: Exemplar events of WS in other species (*D. takahashii, D. kikkawai, D. malerkotliana, D. pseudoananassae, D. willistoni, D. mojavensis* and *D. mercatorum*) during courtship or before copulation.

Table 1: *Drosophila* strains used in this study.

