## Extended Data Figures for "Ancestral neural circuits potentiate the origin of a female sexual behavior"

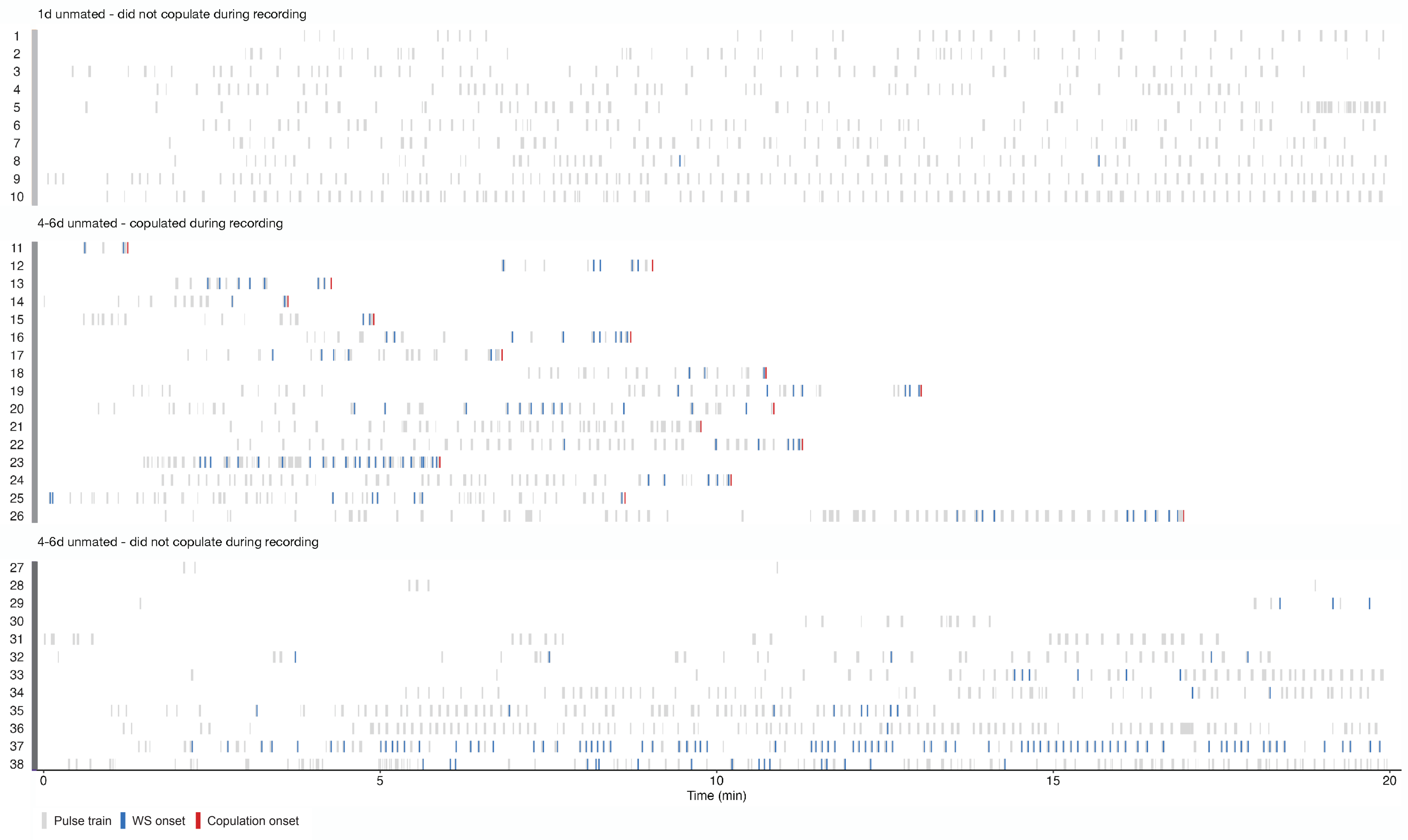


**Extended Data Fig. 1: Full-length behavioral ethograms of *D. santomea* courting pairs, including the ones shown in Fig.1a.**
Each row corresponds to one courting pair, with the numbers to the left representing their pair ID.

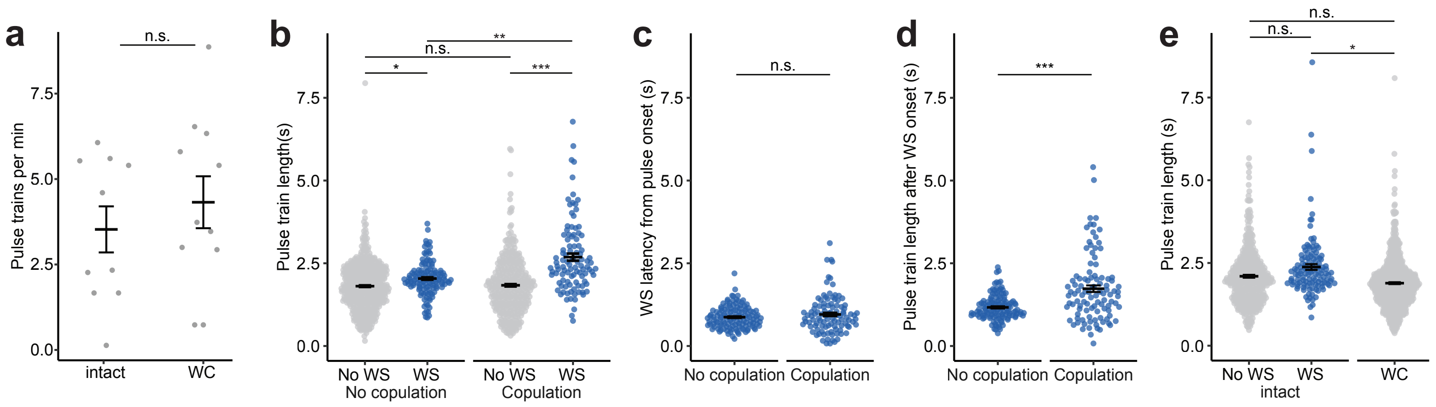


**Extended Data Fig. 2: Additional behavioral analyses linked to Fig. 2 showing WS-dependent modulation of pulse train length.**
**a,** Number of pulse trains per minute that males produced when paired with intact or wing-cut (WC) females. Only data from pairs that did not copulate during the recording period are shown.
**b,** Length of pulse trains separated by whether they elicited WS and whether the pair copulated during the recording period.
**c,d,** Latency of WS from pulse train onset (**c**) and the length of pulse train after WS onset (**d**), respectively, separated by whether the pair copulated during the recording period. In (**d**), the non-parametric Mann-Whitney U test is used to test for statistical difference between the two groups.
**e,** Length of pulse trains in pairs with intact females, separated by whether they elicited WS, and in pairs with WC females. Only data from pairs that did not copulate during the recording period are shown.
Unless otherwise specified, error bars show mean±SEM, and statistical significance was tested with linear mixed models using pair identity as a random effect. *** $p<0.001$, ** $p<0.01$, * $p<0.05$, n.s. not significant.

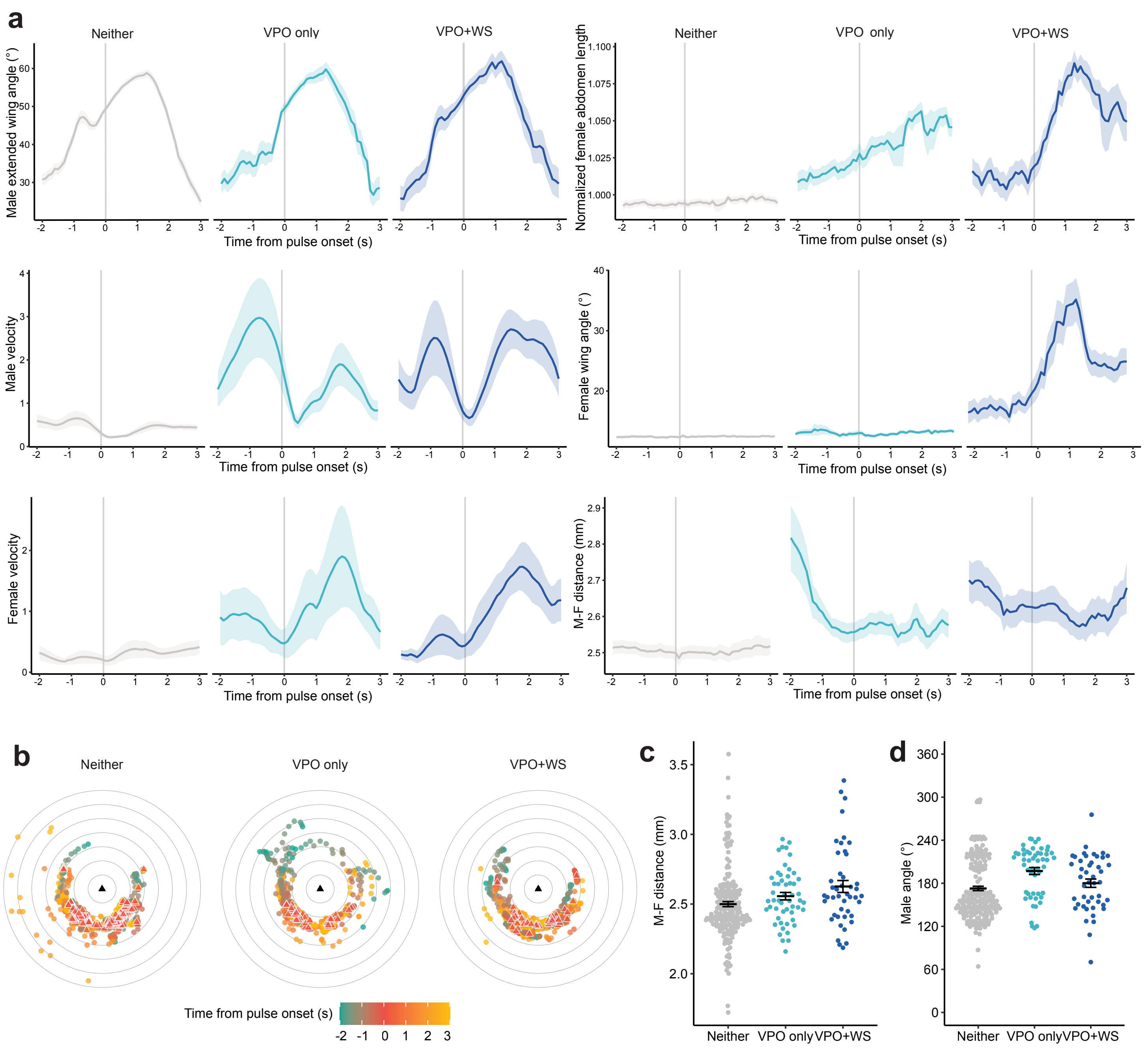


**Extended Data Fig. 3: Female and male behavioral parameters during pulse events.**
**a,** Mean male extended wing angle, male velocity, female velocity, normalized female abdomen length, female wing angle, and distance between male and female thoraces, separated by event type. Pulse onset is marked with a vertical gray line. Shaded areas represent the SEM.
**b,** Male position in female-centered coordinates during all pulse events, separated by event type. The female is represented by a triangle in the center, with the head pointing up. Each ring represents 1 mm. Male position at pulse onset is marked as a red triangle.
**c,** Distance between male and female thoraces at pulse onset compared across all event types.
**d,** Male angle relative to the female body axis compared across all event types. At 180°, the male is directly behind the female.
Error bars show mean±SEM.

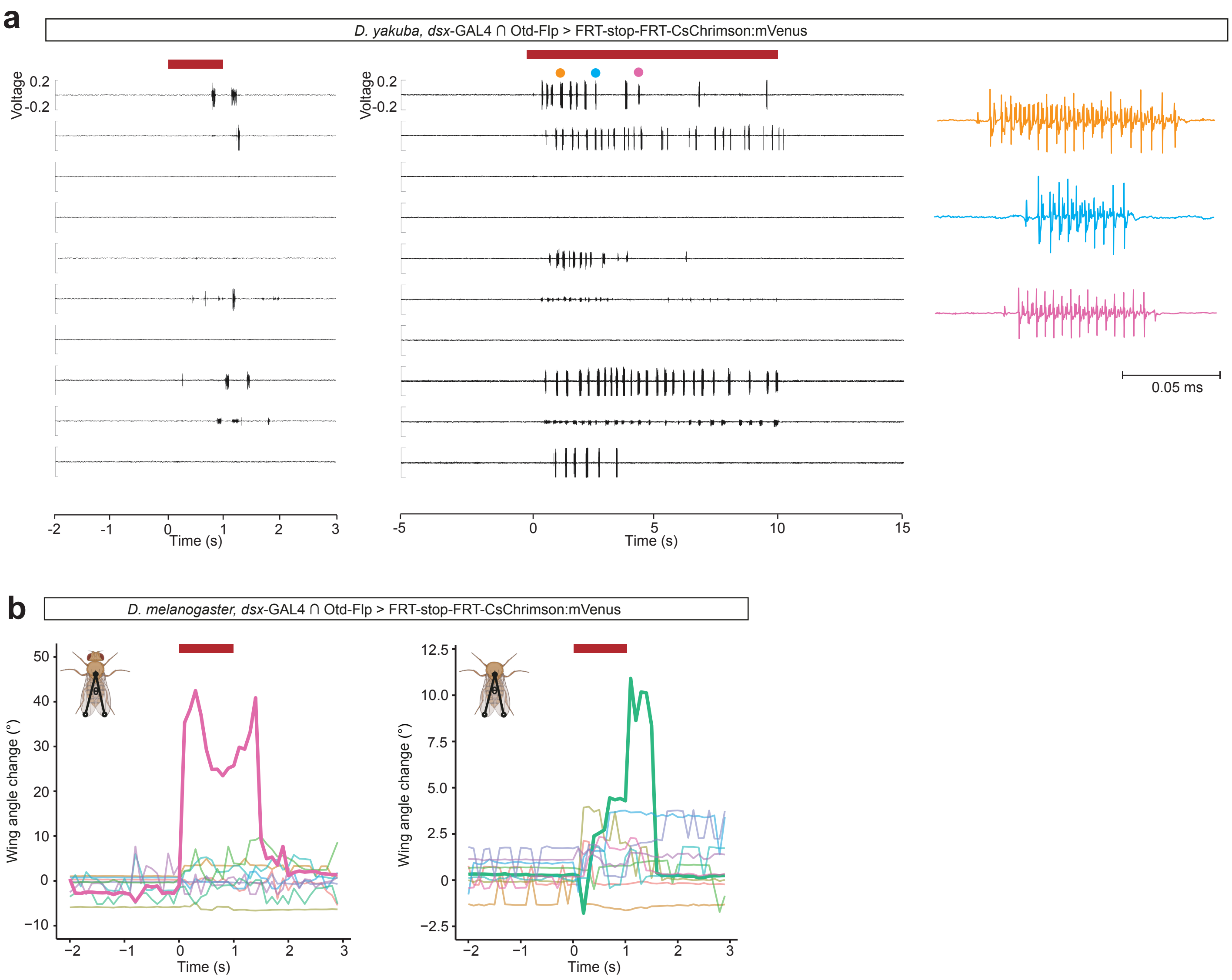


**Extended Data Fig. 4: Additional behavioral phenotypes of activating *dsx* neurons in *D. yakuba* and *D. melanogaster*.**
**a,** Audio traces upon activating *dsx* brain neurons in decapitated *D. yakuba* females using a 1s (left) or a 10s (right) activation scheme with a light intensity of of 4 μW/mm². Each row represents one individual. Three examples of female song events (right) show the production of polycyclic signals with stereotypic waveforms in variable lengths.
**b,** Wing angle change in intact *D. melanogaster* females at 1.6 μW/mm², and decapitated females at 0.8 μW/mm². Activation window is denoted by red bars. Each line with a different color represents one individual. Individuals showing WS response in this activation window are highlighted with thicker lines.

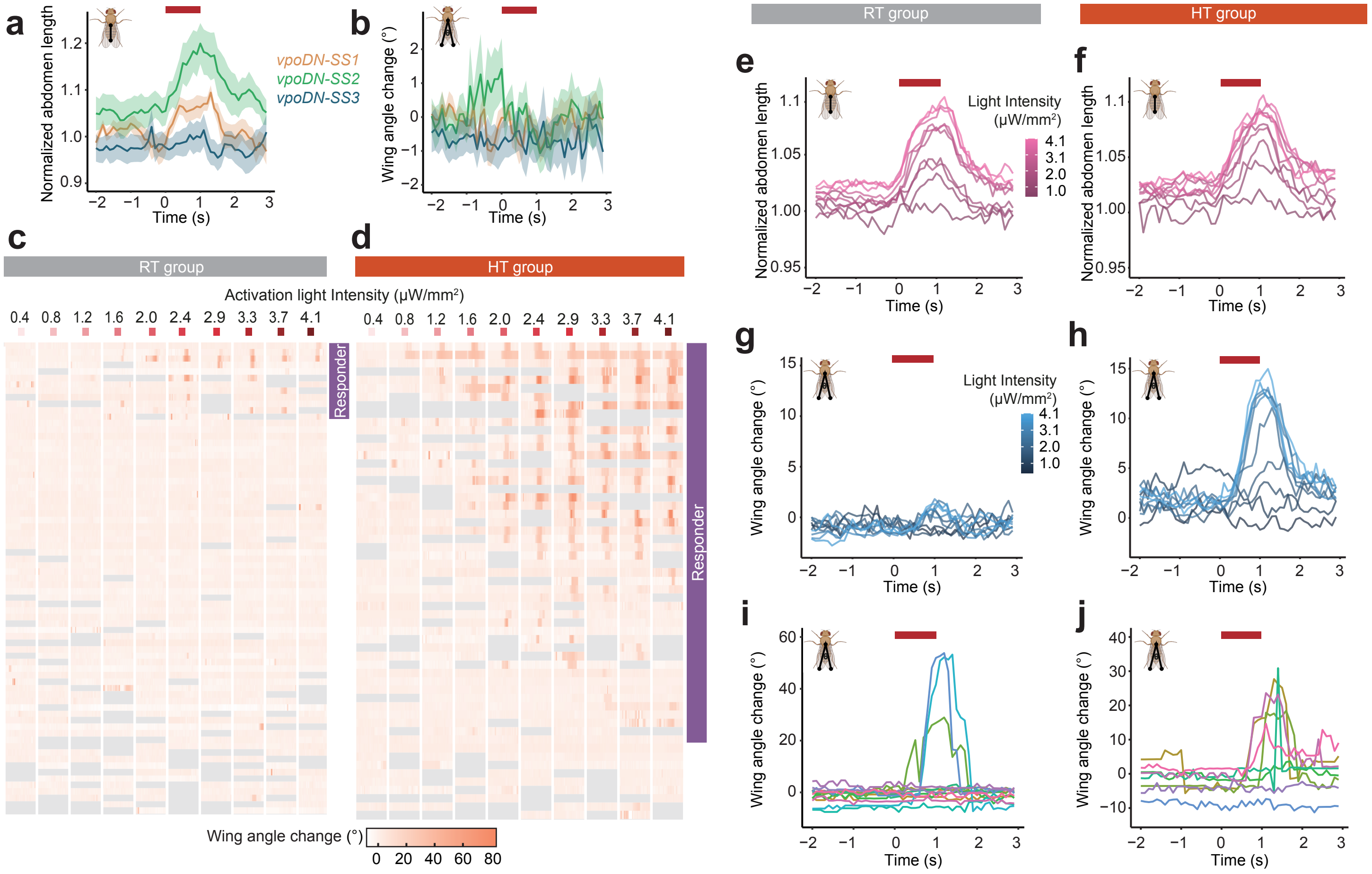


**Extended Data Fig. 5: Additional behavioral analyses associated with Fig. 5 showing developmental temperature-dependent modulation of WS phenotype upon vpoDN activation.**

**a,b,** Mean normalized abdominal length (**a**) and wing angle change (**b**) of *D. melanogaster* vpoDN-SS1, vpoDN-SS2 or vpoDN-SS3 > UAS-CsChrimson:mVenus females at 4.1 μW/mm². Activation window is denoted by red bars. Shaded areas represent the SEM. Inset diagrams illustrate how abdomen lengths or wing angles were measured.

**c,d,** Wing angle changes of *D. melanogaster* vpoDN-SS2 > UAS-CsChrimson:mVenus females from RT group (**c**) and HT group (**d**) across 10 activation bouts with intensity from 0.4 to 4.1 μW/mm². Each row represents one individual. Each column represents a 5-second period centered at 1-second activation window (red bar) with light intensity listed above. Frames with low tracking quality are shown in gray. The manually scored responders are noted on the right side.

**e-h,** Mean normalized abdominal lengths (**e**,**f**) and wing angles (**g**,**h**) of *D. melanogaster* vpoDN-SS2 > UAS-CsChrimson:mVenus females from RT group (**e**,**g**) and HT group (**f**,**h**) at different activation intensities denoted by colors.

**i,j,** Wing angle changes of *D. melanogaster* vpoDN-SS2 > UAS-CsChrimson:mVenus females from RT group (**i**) and HT group (**j**) at 3.3 μW/mm². Ten individuals were randomly selected in the plot and each line represents one individual.
