## Supplementary Table for "Ancestral neural circuits potentiate the origin of a female sexual behavior"

| Figure | Species | Strains |
|---|---|---|
| Figs. 1a-f, 2, 3, 6<br>Extended Data Figs. 1-3 | *D. santomea* | STO4 |
| Fig. 1g | *D. santomea* | 18CAR05-10 |
| Fig. 1g | *D. santomea* | 18TRNE01-05 |
| Fig. 1g | *D. santomea* | Q1-02 |
| Figs. 2a, 6 | *D. yakuba* | 14021-0261.02 |
| Fig. 1g | *D. yakuba* | 3-41 |
| Fig. 1g | *D. yakuba* | 4-72 |
| Fig. 1g | *D. yakuba* | 1-03 |
| Fig. 1g | *D. yakuba* | 2-0F11 |
| Figs. 1g, 2a, 5h,i, 6 | *D.melanogaster* | Canton S |
| Figs. 1g, 6 | *D. simulans* | sim5 |
| Figs. 1g, 6 | *D. teissieri* | 14021-0257.01 |
| Figs. 1g, 6 | *D. erecta* | 14021-0224.01 |
| Fig. 4a-i | *D. santomea* | sanw; ; Otd-Flp (2253), *dsx*-GAL4/FRT-stop-FRT-CsChrimson:mVenus (2253) |
| Fig. 4a-i<br>Extended Data Fig. 4a,b | *D.melanogaster* | w1118; Otd-Flp (attP40)/+; dsx-GAL4/FRT-stop-FRT-CsChrimson:mVenus (vk5) |
| Fig. 4a-i<br>Extended Data Fig. 4c,d | *D. yakuba* | yakw; FRT-stop-FRT-CsChrimson:mVenus (2180)/+; *dsx*-GAL4/Otd-Flp (2285) |
| Fig. 5a-h<br>Extended Data Fig. 5a-j | *D.melanogaster* | vpoDN-SS2: 20xUAS-CsChrimson-mVenus (attP18)/w1118; 45670-AD(p40)/Cyo; 52F12-DBD(p2)=ss50795 |
| Extended Data Fig. 5a,b | *D.melanogaster* | vpoDN-SS1: 20xUAS-CsChrimson-mVenus (attP18)/w1118; 31D07-AD(p40)/Cyo; 52F12-DBD(p2)=ss50200 |
| Extended Data fig. 5a,b | *D.melanogaster* | vpoDN-SS3: 20xUAS-CsChrimson-mVenus (attP18)/w1118;45670-AD(p40)/Cyo;10A09-DBD(p2)=ss53451 |
| Fig. 6 | *D. eugracilis* | 14026-0451.02 |
| Fig. 6 | *D. lucipennis* | unknown |
| Fig. 6 | *D. biarmipes* | G#224 |
| Fig. 6 | *D. takahashii* | unknown |
| Fig. 6 | *D. elegans* | unknown |
| Fig. 6 | *D. gunungola* | sk |
| Fig. 6 | *D. kikkawai* | 14028-05161.14 |
| Fig. 6 | *D. parabipectinata* | VT04-70 Vietnam |
| Fig. 6 | *D. malerkotliana* | C1Z19-L9 |
| Fig. 6 | *D. ananassae* | 14024-0371.13 |
| Fig. 6 | *D. bipectinata* | THT 08 Taiwan |
| Fig. 6 | *D. pseudoananassae* | R186 Puerto Princessa |
| Fig. 6 | *D. pseudoobscura* | 14022-0121.94 |
| Fig. 6 | *D. willistoni* | 14030-0814.24 |
| Fig. 6 | *D. ezoana* | 15010-0971.00 |
| Fig. 6 | *D. mojavensis wrigleyi* | 15081-1352.22 |
| Fig. 6 | *D. mercatorum* | unknown |
